# Sensitivity analysis of agent-based simulation utilizing massively parallel computation and interactive data visualization

**DOI:** 10.1101/510057

**Authors:** Atsushi Niida, Takanori Hasegawa, Satoru Miyano

## Abstract

An essential step in the analysis of agent-based simulation is sensitivity analysis, which namely examines the dependency of parameter values on simulation results. Although a number of approaches have been proposed for sensitivity analysis, they still have limitations in exhaustivity and interpretability. In this study, we propose a novel methodology for sensitivity analysis of agent-based simulation, MASSIVE (Massively parallel Agent-based Simulations and Subsequent Interactive Visualization-based Exploration). MASSIVE takes a unique paradigm, which is completely different from those of sensitivity analysis methods developed so far, By combining massively parallel computation and interactive data visualization, MASSIVE enables us to inspect a broad parameter space intuitively. We demonstrated the utility of MASSIVE by its application to cancer evolution simulation, which successfully identified conditions that generate heterogeneous tumors. We believe that our approach would be a de facto standard for sensitivity analysis of agent-based simulation in an era of ever-growing computational technology. All the result form our MASSIVE analysis is available at https://www.hgc.jp/~niiyan/massive.

## Introduction

Agent-based simulation is a useful tool to address questions regarding real-world phenomena and mechanisms and widely employed in the natural sciences and engineering disciplines as well as in the social sciences [1, 2]. An agent-based model assumes autonomous system components called agents and defines rules that specify behaviors of the agents as well as interactions between the agents, and between the agents and environments. One of the major difficulties in agent-based modeling is determining the values of system parameters, which controls the agent behaviors and interactions. Except for simple physical systems where precise values of the system’s parameters are available, it is often the case that estimated parameter values are used for simulation. In such cases, sensitivity analysis is mandatory; namely, we need to perform simulations with various parameter settings to confirm the robustness of the conclusion that was obtained based on the estimated parameter values. Moreover, sensitivity analysis could provide insights into the modeled system as well as identify parameters that are critical for the system dynamics.

So far, a number of approaches have been proposed for sensitivity analysis of agent-based simulation [3]. For example, one-factor-at-a-time (OFAT) sensitivity analysis selects a base parameter setting and varies a target parameter at a time while keeping all other parameters fixed [4]. We then plot the relationship between the target parameter and a summary static to examine the dependency of the summary statistic on the target parameter. However, since an agent-based model generally involves nonlinear interactions between agents and enviroments, it is desirable to examine multiple combinations of parameters in sensitivity analysis. Global sensitivity analysis aims to address this point by sampling a summary statistic over a wide parameter space involving multiple parameters [5]. The sampled summary statistic is fit to parameters by in a similar fashion as regular regression is done, for instance by means of ordinary least squares. Otherwise, we employ Sobol’s method, which estimates the contributions of different combinations of parameters to the variance of the summary statistic while making the assumption that all parameters are independent [6]. However, these global sensitivity analyses still appears to be insufficient to comprehensively grasp how the parameters that were judged to be influential control behaviors of the agent model.

This paper proposed a new approach to sensitivity analysis termed MASSIVE (Massively parallel Agent-based Simulations and Subsequent Interactive Visualization-based Exploration). MASSIVE conquers the limitation in existing methods by taking advantage of two currently rising technologies: massively parallel computation and interactive data visualization. MASSIVE employs a full factorial design involving a multiple number of parameters (i.e, test every combination of candidate values of the multiple parameters), which could broadly cover a target parameter space but needs a huge computational cost. To deal with this problem, we utilized a supercomputer, in which agent-based simulations with different parameter settings and the following post-processing step of simulation results are parallelly performed. The massively parallel simulations generate massive results, which then poses a problem for interpretation. This problem was solved by developing a web-based tool that interactively visualizes not only values of multiple summary statistics but also results from simulations with each parameter settings. MASSIVE realizes comprehensive sensitivity analysis targeting four parameters at once, and we show the utility by analyzing an agent-based model of cancer evolution.

**Figure 1:**
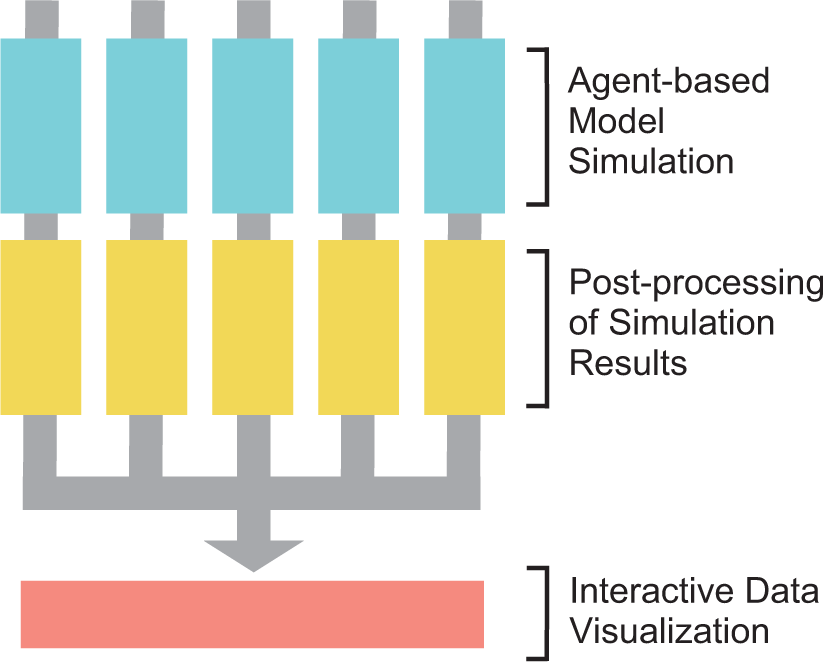
A flow chart of MASSIVE. Agent-based simulations and the following-post processing step are parallelly performed in a supercomputer. Results are then collected and subjected to interactive data visualization.

## Methods

### Agent-based simulation of cancer evolution

Cancer is an evolutionary disease, where a normal cell transforms to a malignant cell population by repeating steps of driver mutation acquisition and subsequent natural selection. Recent genomic studies have demonstrated multiple cell populations that have different genomes were generated during the tumor evolution. This phenomenon is called intratumor heterogeneity and we can use agent-based simulation for understanding mechanisms that generate intratumor heterogeneity [7, 8]. As an example of the application of MASSIVE, we analyze an agent-based model of cancer evolution, where an agent corresponds to each cell in a tumor. The simulation starts from one cell without mutations. In a unit time, a cell divides into two daughter cells with a probability *g* (we assume the cell is immortalized and just divide without dying). In each cell division, each of the two daughter cells acquires *k ∼* Pois(*m/*2) mutations. We assumed mutations acquired by different division events occur different genomic positions and each cell can accumulate *N* mutations at maximum. In this study, we assumed that all mutations are driver mutations which increase the cell division rate. When the cell acquires mutations, the cell division rate increases *f* fold per mutation. In addition to the effect of mutations, we also consider an effect of spatial resource limitation. The simulation is performed on a one-dimensional lattice with free-ends, where a cell divides while pushing out neighboring cells (namely, the model can be regarded as a cellular automaton model). We assumed a resource bias; i.e., resources that are needed for cell division are provided from both the ends of the one-dimensional lattice and are subject to exponential decay. In the presence of the resource bias, the cell division probability for the cell that has *n* (= Σ *k*) mutations and is positioned at the *i*-th site in a one-dimensional population composed of *p* cells (1 ≤ i ≤ *p*) is defined as follows: *g* = *g*_0_ ⋅ *f*^*n*^ ⋅ (2^(*i*−1)*/d*^ + 2^(*p*−*i*)*/d*^)/2 (where *g*_0_ is a base division probability and *d* is a half-distance of the exponential decay). In each time step, every cell is subject to a cell division trial, which is repeated until population size *p* reached at *P* or time reach at *T*.

**Figure 2:**
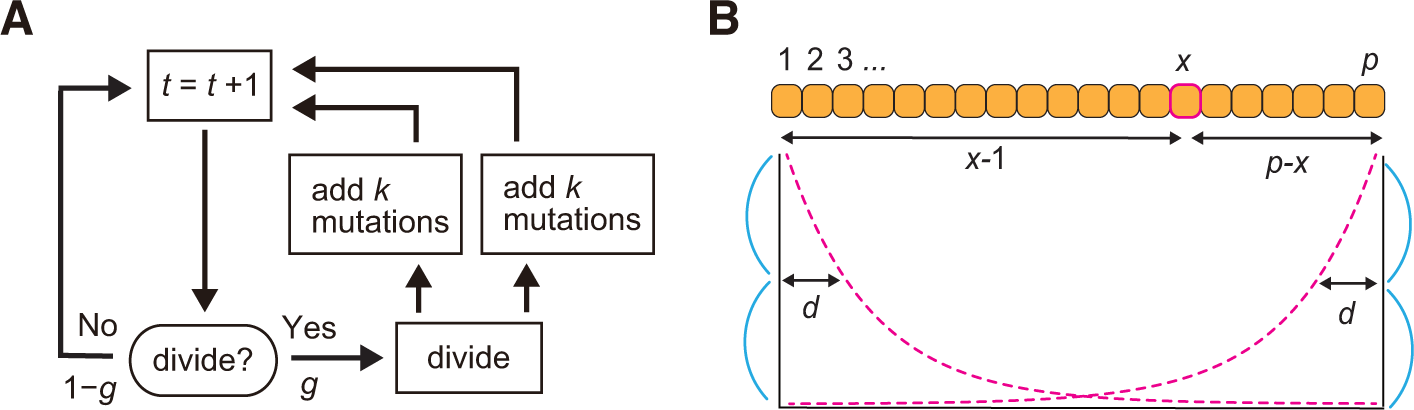
An agent-based model of cancer evolution. (A) A flowchart of the simulation. In each time step, a cell divides into two daughter cells with a probability *g*. In each cell division, each of the two daughter cells acquires *k* ~ Pois(*m/*2) mutations. (B) A growth rule on a one-dimensional lattice. Cell divides while pushing out neighboring cells on a one-dimensional lattice with free-ends. Resources are provided from both the ends and subject to exponential decay. when a cell has *n* (= Σ *k*) mutations and is positioned at the *i*-th site in a one-dimensional population composed of *p* cells (1 ≤ *i ≤ p*), cell division probability, *g* = *g*_0_ ⋅ *f*^*n*^ ⋅ (2^(*i−*1)*/d*^ + 2^(*p−i*)*/d*^)*/*2 (where *g*_0_ is a base division probability and *d* is a half-distance of the exponential decay)

### Post-processing of simulation results

A result from the cancer evolution simulation was evaluated by visualizing mutation profile and calculating summary statistics from the mutation profile. From a simulated tumor, we randomly sampled 1000 cells and obtained their mutation profile, which is represented as a binary matrix whose row and column index mutations and cells, respectively. The mutation profile matrix was visualized as a heat map in Figure 3, where columns were ordered based on the original cell positions and rows were reordered by hierarchical clustering. We attached colored bands indicating mutation types and the distribution of the resources on the left and top of the heat map, respectively. For mutation type, we defined mutations whose frequency is more than 0.95 as clonal mutations and the others as subclonal mutations. To quantitatively evaluate the simulation result, we also calculated a number of summary statistics from the mutation matrix. We obtained several statistics for evaluating intratumor heterogeneity as well as other basic statistics (Table 1). For example, after removing mutations whose frequency less than 0.1 or 0.05, proportions of different subclones (cell subpopulations with different mutations) were obtained and Shannon index *H* was calculated by the following formula: 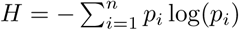, where *n* is the total number of the different subclones and *p*_*i*_ is the proportion of each subclone. Simpson index 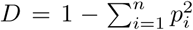 was similarly calculated for each of the two mutation frequency cutoffs.

**Figure 3:**
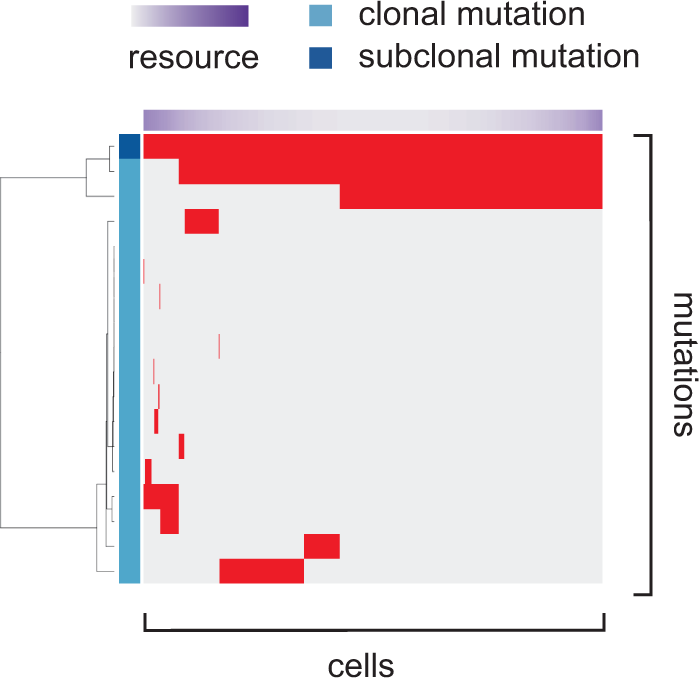
Visualization of the mutation profile matrix. From a simulated tumor, we randomly sampled 1000 cells and their mutation profile is presented as a mutation profile matrix. Columns corresponding to cells were ordered based on the original cell positions and rows corresponding to mutation were reordered by hierarchical clustering. We attached colored bands indicating mutation types and the distribution of the resources on the left and top of the heat map, respectively. For mutation type, we define mutations whose frequency is more than 0.95 as clonal mutations and the others as subclonal mutations.

**Table 1:** Variables

| symbol | description |
| --- | --- |
| $k$ | number of mutations obtained in a cell division |
| $n$ | number of mutations accumulated in a cell |
| $p$ | population size |
| $g$ | cell division probability |
| $i$ | coordinate on the one-dimensional lattice |

### Massively parallel simulation

To cover a sufficiently large parameter space in the sensitivity analysis, we employed a supercomputer, SHIROKANE4 (at human genome center, IMSUT). The simulation and post-processing steps for different parameter settings were parallelized on Univa Grid Engine. We employed a full factorial design involving four parameters, *m*, *f*, *P* and *d*, while other parameters were fixed. The values of all parameters are listed in Table 2. For convenience, we converted the parameters as follows: *m^′^* = *−* log_10_(*m*), *f′* = log_10_(*f*), *p^′^* = log_10_(*P*) and *d^′^* = log_10_(*d*). We then tested every combination of *m^′^* ∈ {1, 2, 3}, *f′* ∈ {0.1, 0.2, 0.3*, …*, 1.0}, *p′* ∈ {3, 4, 5} and *d′* ∈ {1, 2, 3, 4, 5}, respectively. For each parameter setting, we performed 50 Monte Carlo trials, which leads to 3 *×* 10 *×* 3 *×* 5 *×* 50 = 22500 simulations in total. On average, each simulation took about 1046 CPU core seconds. Therefore, our study needed 1046 *×* 22500 = 23528298 CPU core seconds *≈* 273 CPU core days in total. However, by parallelizing thousands of simulations, they finished within several hours.

**Table 2:** Parameters

| symbol | description | value |
| --- | --- | --- |
| $m$ | mutation rate | $\{10^{-1}, 10^{-2}, 10^{-3}\}$ |
| $N$ | maximum number of mutations accumulated in a cell | 3 |
| $f$ | cell division probability increase per mutation | $\{10^{0.1}, 10^{0.2}, 10^{0.3}, \dots, 10^{1.0}\}$ |
| $g_0$ | base cell division probability | $10^{-4}$ |
| $d$ | half-distance of the exponential resource decay | $\{10^1, 10^2, 10^3, 10^4, 10^5\}$ |
| $P$ | maximum population size | $\{10^3, 10^4, 10^5\}$ |
| $T$ | maximum number of time steps | $10^6$ |

**Table 3:** Summary statistics

| name | description |
| --- | --- |
| mutation count | number of all mutations |
| clonal mutation count | number of clonal mutations |
| subclonal mutation count | number of subclonal mutations. |
| clonal mutation proportion | proportion of clonal mutation count to mutation count |
| subclonal mutation proportion | proportion of subclonal mutation count to mutation count |
| Shannon index 0.1 | Shannon index calculated with a mutation frequency cutoff of 0.1 |
| Shannon index 0.05 | Shannon index calculated with a mutation frequency cutoff of 0.05 |
| Simpson index 0.1 | Simpson index calculated with a mutation frequency cutoff of 0.1 |
| Simpson index 0.05 | Simpson index calculated with a mutation frequency cutoff of 0.05 |
| time | number of time steps when simulation is finished |
| population size | number of cells when simulation is finished |

### Interactive data visualization

To analyze massive results from massively parallel simulations, we build a new visualization tool termed the MASSIVE viewer. After all the simulations and post-processing steps were finished, the mean of each summary statistic was calculated over 50 Monte Carlo trials for each parameter settings. The data were then prepared as an input of MASSIVE viewer on the web server, together with image data of mutation profile matrix heat maps. The MASSIVE viewer reads the inputs and interactively visualize them on web browser utilizing a javascript library, D3.js [9]. A concept central to the MASSIVE viewer is the multilayer heat map, which was introduced in our previous study [10]. The multilayer heat map visualizes slice matrices of a fourth-order tensor *T* (*i, j, k, l*). A slice matrix contains tensor elements which have specified indices in two of the four orders; namely, when a colon denotes the set of all indices for a particular order, slice matrices are represented as: *T* (:, :*, k, l*), *T* (:*, j,* :*,l*), *T* (:*, j, k,* :) … etc. (we referred to them as *i*-*j*, *i*-*k*, *i*-*l* slice matrices etc.) Since we employed a full factorial design involving four parameters in massively parallel simulations, each of the averaged summary statistics is represented as a fourth-order summary statistics tensor, *S*(*x, y, z, w*), where *x*, *y*, *z* and *w* index values tested for each type of the four parameters.

The MASSIVE viewer has two different modes of visualization: the focused and comparative views. The focused view mode is useful for inspecting the parameter sensitivity of one summary statistic in the whole parameter space tested. The page of the focused view mode is divided into three parts: control, heat map and simulation instance panels. The top is the control panel, which specifies the type of summary statistic to be visualized and which parameters are indexed by each of *x*, *y*, *z* and *w*. Under the control panel, the heat map panel is located. Heat maps at the left part of the heat map panel present all *x*-*y* slice matrices. each cell in the heat maps corresponds to each of the tested parameter sets. At the right part of the heat map panel, the *x*-*y* slice matrix under the mouse cursor at the left part is zoomed out. We can click to highlight a heat map at the left part so that the highlighted one is fixedly displayed at the right part. Under the heat map panel, a simulation instance panel displays 5 mutation profiles together with values of parameters and summary statistics. The mutation profiles were produced from 5 of the 50 Monte Carlo trials with the parameter set corresponding to the heat map cell under the mouse cursor at the heat map panel. We can fix mutation profiles by clicking to highlight a heat map cell at the right part of the heat map panel.

The comparative view mode is used for comparing each summary statistics. Similarly to the focused mode, the page of the comparative view mode consists of control, heat map and simulation instance panels. In the heat map panel, we have three heat maps presenting three *x*-*y* slice matrices of common *z* and *w* indices for three summary statistics. In addition to the parameter and summary statistics setting employed in the focused mode, the control panel in the comparative mode specifies the parameter values tied with the *z* and *w* indices; i.e., we can specify the parameter values that define the third and fourth order indices of the displayed *x*-*y* slice matrices. The control panel also specifies either of the two scale types: the absolute or relative scales. If we select the relative scale, the color scale of each of the three heat maps is prepared for each time the values of the parameters associated with *z* and *w* are changed; namely, one color scale is used for each *x*-*y* slice matrix. If we select the absolute scale, the color scale is unchanged; namely, a common color scale is used for all *x*-*y* slice matrices from one summary statistics tensor. Similarly to the focused mode, the simulation instance panel display mutation profiles, which we can interactively explore across the tested parameter space by hovering over and clicking the heat map cells in the heat map panels.

## Results

In this section, we explain the utility of MASSIVE using the example of the agent-based simulation of cancer evolution. We examined the behavior of the simulation model with different values of four parameters, *f′*, *d′*, *m′* and *p′*. A larger *f′* means a stronger mutation effect, a larger *d′* means a stronger resource bias, a larger *p′* means a larger maximum population size, and a larger *m′* means a smaller mutation rate. First, we examined different types of statistics for quantifying intratumor heterogeneity. As shown by the comparative view mode page (we ask reader to click the hyperlink and explore the result interactively), three different statistics, `Shannon index 0.1, Shannon index 0.05` and `Simpson index 0.1`, shows similar profiles across the whole parameter space. `Simpson index 0.05` also shows a similar pattern. We can confirm that these heterogeneity atatistics well represent actual heterogeneity by checking mutation profiles in the simulation instance panel. Based on these observations, we hereafter focused on `Shannon index 0.1` as a representative statistic for a measure of intratumor heterogeneity.

**Figure 4:**
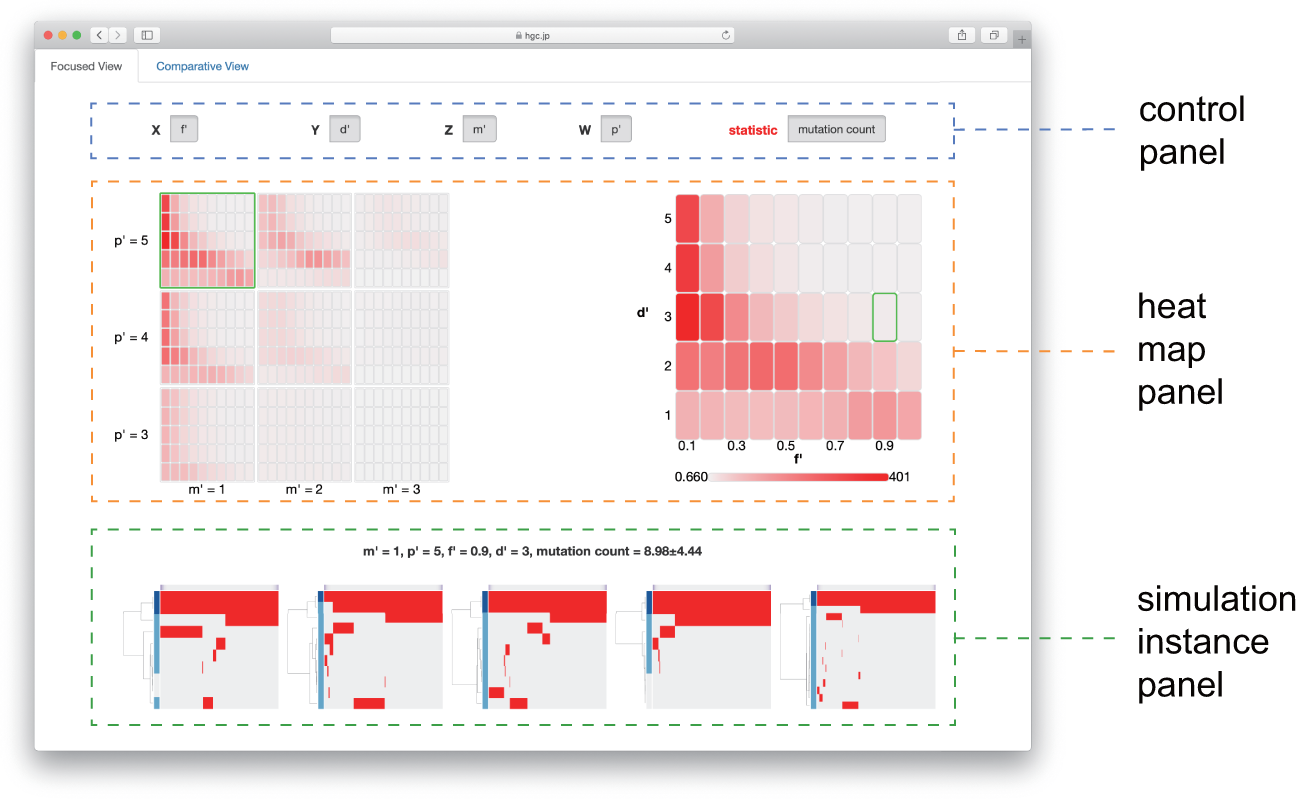
Focused view mode of the MASSIVE viewer. The page of the focused view mode consists of control, heat map and simulation instance panels. The control panel specifies visualization settings, the heat map panels presents heat maps for a selected statistics, and the simulation instance panel presents 5 mutation profiles from the parameter set specified by the position of the mouse cursor on the heat maps.

**Figure 5:**
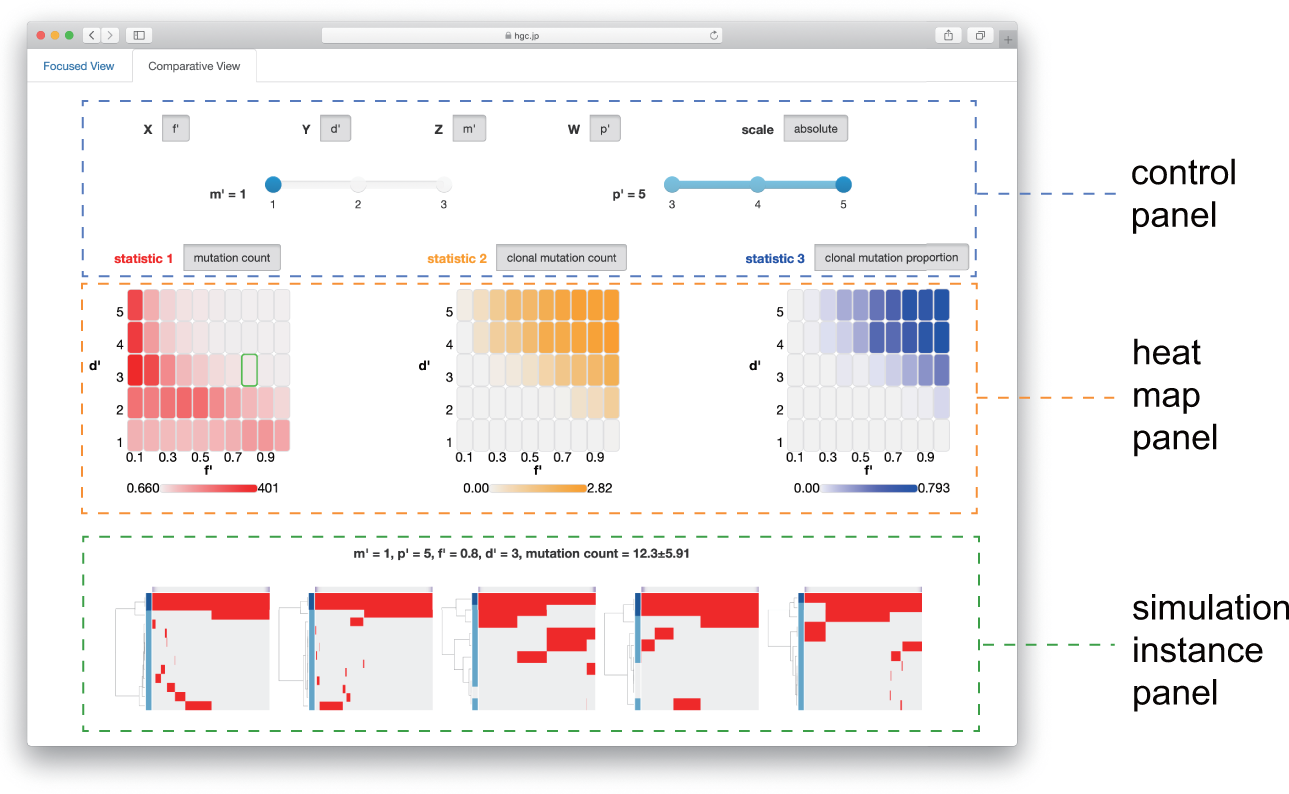
Comparative view mode of the MASSIVE viewer. Similarly to the focused mode, the page of the comparative view mode consists of control, heat map and simulation instance panels. Differently from the focused mode, the heat map panels presents heat maps for three selected statistics.

As shown by the focused view mode page, the heterogeneity measure, Shannon index 0.1 is a complex function of *f′*, *d′*, *m′* and *p′*. As expected, a decrease of *m′* and an increase of *p′* generally leads to an increase of the heterogeneity measure; i.e., the heterogeneity is enhanced as the tumor grows and the mutation rate increases. When we focus on a parameter plain of *p′* = 5 and *m′* = 1, less heterogeneity is observed at high *f′* and high *d′* (i.e., with strong mutations and with less resource bias). This observation is interpretable as follows. When no resource bias and strong driver genes are assumed, the clone acquiring the first driver mutation rapidly expand to serially obtain more driver mutations, which leads to a homogeneous tumor. On the other hand, a resource bias prompts generation of independent clones at both sides on the one-dimensional lattice, which leads to a heterogeneous tumor. With weak driver genes, before the clone that acquired the first driver mutation becomes dominant, other clones that acquired different mutations expand, which leads to a heterogeneous tumor even without a resource bias. Collectively, our exemplifying analysis successfully provided insights into heterogenous cancer evolution, demonstrating the utility of the MASSIVE analysis.

## Discussion

In this study, we introduced a novel methodology for sensitivity analysis of agent-based simulation, MASSIVE. MASSIVE takes a unique paradigm, which is completely different from those of sensitivity analysis methods developed so far; by combining massively parallel computation and interactive data visualization, MASSIVE enables us to inspect a broad parameter space intuitively. We demonstrated the utility of MASSIVE by its application to cancer evolution simulation, which successfully identified conditions that generate heterogeneous tumors.

However, the current version of MASSIVE still has a major limitation; it targets at most a four-dimensional parameter space. To analyze a higher dimensional parameter space, methodological improvements are necessary for both massively parallel computation and interactive data visualization. In addition to the hardware advancement, it is necessary to develop an approach for efficient sampling of the huge parameter space in order to enable the massively parallel simulation that can overcome the curse of dimensionality. Results from simulation trials in the higher dimensional space are also not easily interpretable even by employing interactive data visualization. To conquer this limitation, the combination with dimension reduction techniques such as autoencoder [11] might be useful.

Parameter estimation by comparing simulation trials with real data is also an essential step in simulation analysis. both need a large number of simulation trials and it is conceivable that results of parameter estimation are performed simultaneously with sensitivity analysis. In our framework, such a fusion of sensitivity analysis and parameter estimation can be easily implemented; the parameter subspace that generates simulation results similar to the real data can be presented in the visualization step. Especially, Approximate Bayesian Computation (ABC) is recently emerging as a popular tool for parameter estimation [12, 13]. Although ABC requires a large number of simulations trials compared to conventional parameter estimation methods, it can evaluate the confidence of estimated parameter values by computing the posterior distributions of the parameter values. These features of ABC also appear to have a strong affinity with our framework.

Collectively, this work proposes a novel approach for sensitive analysis of agent-based simulation based on the combination of massively parallel computation and interactive data visualization. We believe our approach is a de facto standard for sensitivity analysis of agent-based simulation in an era of evergrowing computational technology.

